# Genomic dissection of the microevolution of Australian epidemic *Bordetella pertussis*

**DOI:** 10.1101/2022.01.05.475016

**Authors:** Zheng Xu, Dalong Hu, Laurence Don Wai Luu, Sophie Octavia, Anthony D Keil, Vitali Sintchenko, Mark M. Tanaka, Frits R. Mooi, Jenny Robson, Ruiting Lan

## Abstract

Whooping cough (pertussis) is a highly contagious respiratory disease caused by the bacterium *Bordetella pertussis*. Despite high vaccine coverage, pertussis has re-emerged in many countries and caused two large epidemics in Australia since 2007. Here, we undertook a genomic and phylogeographic study of 385 Australian *B. pertussis* isolates collected from 2008 to 2017. The Australian *B. pertussis* population was found to be composed of mostly *ptxP3* strains carrying different *fim3* alleles, with *ptxP3-fim3A* genotype expanded far more than *ptxP3-fim3B*. Within the former, there were six co-circulating epidemic lineages (EL1 to EL6). The multiple ELs emerged, expanded, and then declined at different time points over the two epidemics, likely driven by immune selection from pertussis vaccination and natural infection in addition to local and global transmission events. Both hard and soft selective sweeps through vaccine selection pressures determined the current *B. pertussis* population dynamics. Relative risk analysis found that once a new *B. pertussis* lineage emerged, it was more likely to spread locally within the first 1.5 years. However, after 1.5 years, any new lineage was likely to expand to a wider region and became no longer spatially structured across the country. Phylogenetic analysis revealed the expansion of *ptxP3* strains was also associated with replacement of the type III secretion system allele *bscI1* with *bscI3*. This study advanced our understanding of the epidemic population structure and spatial and temporal dynamics of *B. pertussis* in a highly immunised population.

## Introduction

*Bordetella pertussis* is the causative agent of the contagious respiratory disease, whooping cough [1]. Although the introduction of whole-cell vaccines (WCVs) has achieved a successful reduction in *B. pertussis* incidence, pertussis re-emerged in the late 1990s in many developed countries after the acellular vaccines (ACVs) replaced the WCV from 1999 [2-4]. Based on the World Health Organisation (WHO) estimates, there were 24.1 million pertussis cases in 2014 with 160,700 deaths in children less than 5-year-old, despite around 86% immunisation coverage with pertussis vaccines globally [5]. In Australia, pertussis epidemics have been occurring cyclically every three to five years. The largest pertussis epidemic started in 2007 and peaked with nearly 40,000 cases reported in 2011 [6], followed by a smaller epidemic from 2013 to 2017.

Several factors are thought to have contributed to the re-emergence of pertussis. The adaptation to vaccine-induced immunity is believed to be an important factor for pertussis re-emergence since genotypic and phenotypic divergences between vaccine strains and circulating strains of *B. pertussis*, especially with respect to genes encoding vaccine antigens, have been reported globally [4,7-11]. For example, pertactin (Prn) is one of the antigens included in acellular pertussis vaccines and strains which do not produce pertactin have emerged recently and became predominant in some countries [12,13]. A mouse infection study compared a Prn-negative isolate with a Prn-positive wild type isolate and showed that the former was more competitive in colonising ACV vaccinated mice [14]. Another example of vaccine-induced genetic adaption is the emergence and spread of strains carrying the *ptxP3* allele across the globe in recent years [4,8]. The *ptxP3* isolates were found to produce a higher amount of pertussis toxin than *ptxP1* isolates, suggesting higher virulence in the *ptxP3* isolates [8]. Higher virulence in *B. pertussis* isolates may have evolved under the selective pressure from vaccines, similar to that observed in *Haemophilus influenzae* Serotype b strains [15].

There are two main ACVs consisting of three and five components [16]. The three component ACV consisted of Prn, pertussis toxin and filamentous haemagglutinin whereas the five component ACV additionally contained fimbriae Fim2 and Fim3. Australian predominantly uses the three component ACV [6]. Variations in ACV antigen genes apart from two key changes discussed above have been observed and has been associated with vaccine selection [6]. For the two fimbrial subunits, Fim2 and Fim3, they are subject to phase variation [17]. Most *B. pertussis* strains produce either Fim2, Fim3 or both. Fim serotypes have changed over time globally and the changes may or may not be vaccine-driven [18]. Dissecting the selection pressures and evolutionary forces in the genetic and phenotypic changes in vaccine antigen genes is often difficult.

Variations in vaccine-related genes also reflected the changes in the population structure of *B. pertussis*. Comparative genomic studies of *B. pertussis* have been conducted to investigate the evolutionary changes contributing to their resurgence. Genome-wide single nucleotide polymorphisms (SNPs) typing was able to divide over 300 global *B. pertussis* isolates into six SNPs clusters (Cluster I to VI) [7]. This, and other, SNP typing study provided phylogenetic evidence supporting the hypothesis that *B. pertussis* has evolved under vaccine selective pressure [4,7]. Subsequently, the emergence of the Cluster I lineage was found to be associated with the emergence of the *ptxP3* allele [9]. Genomic sequencing and phylogenetic analysis of isolates from the 2008-2012 Australian epidemic further defined five epidemic lineages (ELs) within Cluster I in Australia and the parallel evolution of Prn deficiency in different co-circulating lineages [19]. Prn-negative isolates from four of these five ELs continued to circulate in the 2013-2017 Australian pertussis epidemic [13]. Although genetic variation in these isolates has been studied, the temporal and geospatial dynamics of *B. pertussis* population structure changes and evolutionary forces behind have not been elucidated yet.

In addition to vaccine selection, comparative proteomic studies between a Cluster I and a Cluster II strain identified decreased secretion of the type III secretion system (T3SS) in the Cluster I strain [20-22]. This decreased secretion was hypothesised to be associated with a mutation in *bscI*, which encodes the T3SS basal body protein. A mouse infection study with Cluster I and Cluster II strains showed that Cluster I was fitter in both ACV-immunised and unimmunised mice suggesting additional selection pressures [23].

In this study, we used a large set of *B. pertussis* isolates to elucidate the population structure of the pathogen and the factors influencing its evolution during the epidemics in Australia. The fine-scale phylogenetic analyses were integrated with temporal and spatial metadata to examine how *B. pertussis* populations varied over time and space.

## Materials and methods

### Bacterial isolates and genomic DNA preparation

The 385 *B. pertussis* isolates used in this study were sampled from pathology laboratories servicing patients across different states in Australia. Among them, five isolates from 1997-2002 were included to represent strains circulating before the recent epidemics, 302 isolates were sampled from the 2008-2012 epidemic across five states: New South Wales (NSW), Western Australia (WA), Queensland (QLD), Victoria (VIC) and South Australia (SA) and 78 isolates were collected from the 2013-2017 epidemic from two states, WA and NSW, as the other states ceased culturing *B. pertussis* (**Supplementary Table S1**). Due to the poor sequence quality of four isolates from a previous study [19]. these isolates were re-sequenced in this study. Overall, a total of 284 isolates were sequenced in this study. The number of isolates from different states did not reflect the total pertussis incidence of the states. WA has a much smaller population than NSW but more isolates were collected from WA. All available clinical isolates from 2008 to 2017 were included. Annual rates of pertussis notifications in Australia were obtained from the National Notifiable Diseases Surveillance System. Available from: http://www9.health.gov.au/cda/source/cda-index.cfm

Isolates were revived from cryovials stored in freezers at -80°C and inoculated on Bordet-Gengou agar (Becton Dickinson, Sparks, MD, USA, supplemented with 7% horse blood) and aerobically incubated at 37°C for 4-5 days. Pure colonies were harvested for genomic DNA extraction using the phenol-chloroform method [24].

### Genome sequencing and data accessibility

Sequencing libraries were constructed using Nextera XT kit (Illumina). The fragment size distribution of the libraries was analysed using LabChip GX high Sensitivity DNA assay kit (Caliper Life Science, Hopkinton, MA). Whole-genome sequencing (WGS) was performed using Illumina NextSeq (2 × 150 bp) and MiSeq (2 × 300 bp) platform. Raw reads have been submitted to the NCBI GenBank database under the BioProject accession number PRJNA562796. All isolates were sequenced at the Marshall Centre for Infectious Diseases Research and Training (WA, Australia).

### Detection of SNPs and phylogenetic analysis

*De novo* assembly was performed using SPAdes (version 3.14.1) [25]. SNPs were identified using a combination of mapping to the *B. pertussis* reference strain Tohama I (NC_002929.2) using Burrow-Wheeler Alignment (BWA) tool (version 0.7.12) [26], SAMtools (version 0.1.19) [27] and alignment of SPAdes assembled genome to reference Tohama I using progressiveMauve (version snapshot_2015_02-25) [28]. Only SNPs identified consistently by both contig-alignment and mapping methods were used for the downstream analyses. Phylogenetic trees were constructed using RAxML (version 8.2.12) [29] with the complete genome of *B. pertussis* Tohama I as reference and outgroup, applying parameters -m GTRGAMMA and 1,000 bootstrap samples. Determination of IS*481* and IS*1002* insertions was performed using a custom script as previously described in Safarchi *et al*. [19] and Octavia *et al*. [7].

The phylogenetic tree of international *B. pertussis* strains was constructed using 1,452 available *B. pertussis* genomes to locate the spread of ELs. The phylogenetic tree was constructed using Quicktree pipeline (version 1.1) with default settings, in which -gtr -gamma model was used for tree construction with 100 times bootstrap [30].

### BEAST analysis

Divergence time of the nodes on the tree was estimated using the BEAST package (version 1.10.4). Considering that the sampling and sequencing of isolates may not cover all epidemic feathers for the whole outbreak in Australia, a fixed evolutionary model chosen for BEAST may not reflect the truth. However, a comprehensive modelling for estimating the population structure of *B. pertussis* epidemic using sampled genomes is not in the scope of this study. Thus, the model was chosen by statistical evaluation using only the sequenced data, following the strategy described in [31], in which a total number of 24 models (four molecular clock models and six population size models) were set up and run independently using the same data with same MCMC parameters (1,000,000 chains excluding the first 10,000 chains from the posterior distribution as burn-in), and then Effective Sample Size (ESS) value of each model was calculated using Tracer (version 1.7.1). The strict clock with constant population size model was found to have the highest ESS value (ESS=339, the only one model combination with ESS over 200) and was thus selected. The final tree with divergence dates was annotated by TreeAnnotator within the BEAST package.

### Lineage through time (LTT)

The lineage through time (LTT) was performed following Reznick *et al*. [32]. Briefly, LTT refers to the total number of branches at a certain time point from the root to the top of the lineage. In this study, the divergence time of the nodes on the tree was extracted from the BEAST estimation as in **Figure 1**. For a specific lineage, logarithm values of total number of branches were calculated at time points from the root to the top of the lineage. LTT curves were then generated using a custom script. The calculation was conducted phytools package in R (version 3.5.1)[33].

**Figure 1.**
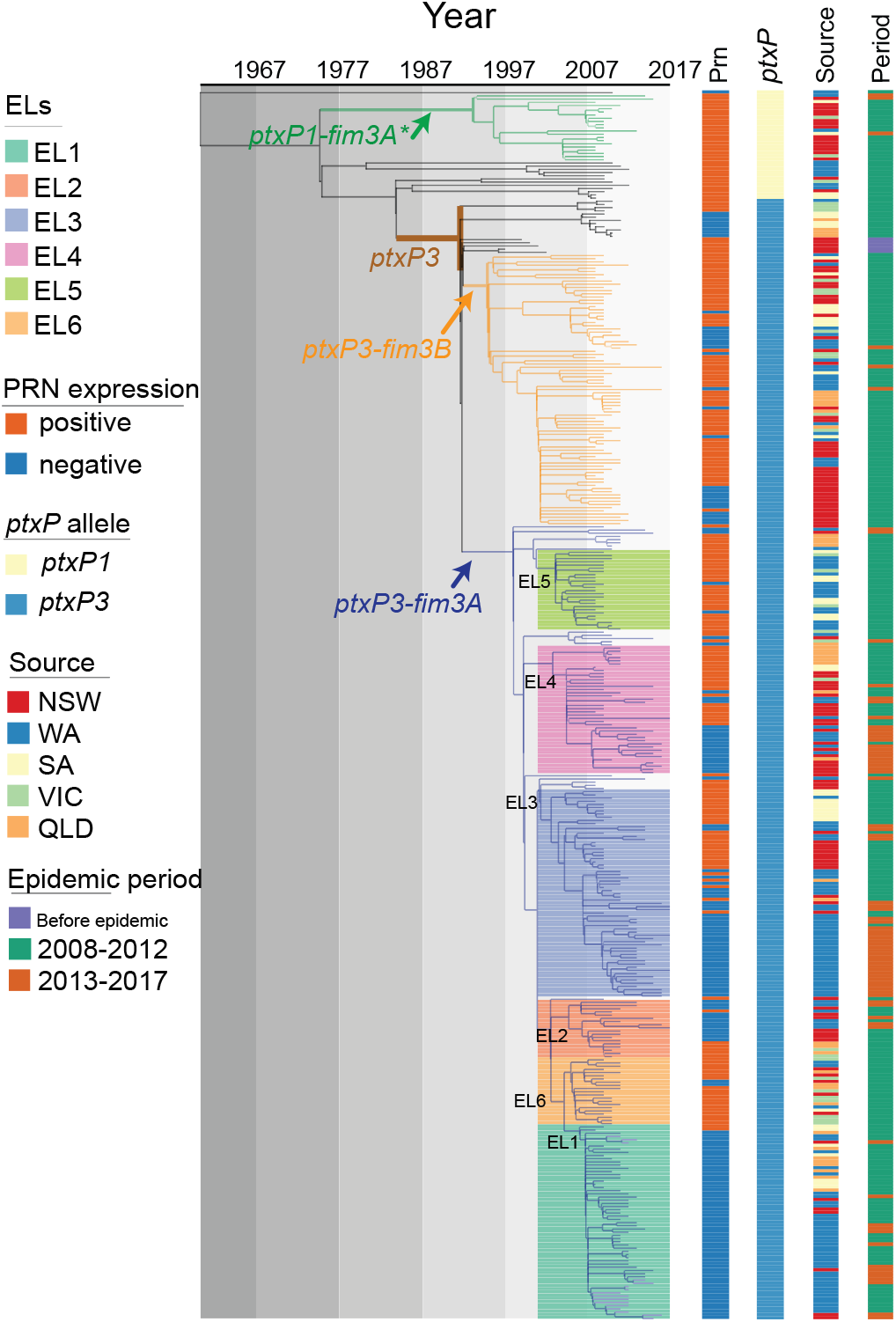
Phylogenetic relationship of Australian epidemic lineages (ELs). A phylogenetic tree of 385 *B. pertussis* isolates was constructed using maximum likelihood method. The branch length was scaled by divergence time as generated by BEAST. The tree was rooted using the *B. pertussis* reference genome Tohama I as an outgroup. Three major lineages were coloured at the branch. The coloured lines and columns from left to right represent ELs, Prn presence or absence, *ptxP* allele types, *fim3* allele types, the geographic source of isolates and period of isolation.

### Relative risk assessment within and between jurisdictions

First, probability was estimated for a pair of cases in geographically defined areas is within the same cluster of transmissions. Then relative risk was computed for the disease transmission as a function of geographic proximity, by comparing probabilities restricted in different ways as the same method shown in [34]. Specifically, the analysis looks at relative risk of infection at different spatial scales within a state and between states.

The calculation of relative risk is based on the following notations. Define *D*_*ij*_ to be the geographic distance between isolates *i* and *j* and let *s*_*i*_ and *s*_*j*_ be the states associated with the isolates. Let *T*_*ij*_ be the estimated time of the most recent common ancestor of isolates *i* and *j*. Let *n* be the total number of isolates being considered. The probability that a pair of geographically defined cases is within the same genetic cluster according to the specified time band is estimated with

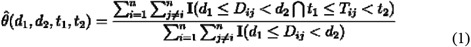

where the sum 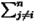 is over the values of *j* between *i* and *n* except where *i = j*, and where **I**(*x*) is the indicator function which evaluates to 1 if *x* is true and 0 otherwise. The relative risk is then estimated by

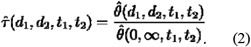

Based on the monthly number of pertussis notifications in Australia from January 1991 until December 2018 from the NNDSS (monthly pertussis notifications were standardised to a 30-day month [35]), four cycle length interval (1.5 years, 3.5 years, 8 years and 16 years) were generated by using the Fast Fourier Transform method [35]. The geographical distance between isolates were mainly divided into five ranges: 0-1 km (same postcode area), 1-20 km (the Perth city area), 20-100 km (average distance to surrounding suburbs of Perth city), 100-260 km (cover the most of suburbs nearby) and >260 km. The > 260 km distance was used as the reference point, because very few pairs of isolates fell into this range of distance in WA. The probability that a pair of interstate isolates, of which one belongs to state *s*, are genetically related under the *t*_1_, *t*_2_ time range for the time to most recent common ancestor (tMRCA) is

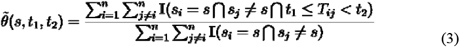

which considers all pairs of isolates in which one is in state *s* and the other is outside *s*. For the relative risk the reference point is the probability that any two interstate pair of isolates is closely related:

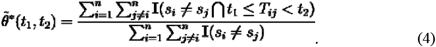

Thus, the relative risk (of interstate transmission for a particular state compared to all states) is

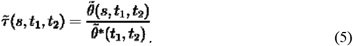

To estimate the uncertainty of the relative risk, 100 times bootstrap resampling were done for each calculation to get the 95% CI intervals. For each time bootstrap process, 385 times independent and random samplings of isolates were performed from the pool of all the 385 isolates in the analysis to construct a bootstrap sample set for the relative risk calculation. All epidemiological factors were preserved in each resampled isolate.

## Results

### Phylogenetic lineages of *B. pertussis* circulating in Australia

All 302 isolates collected in Australia between 2008 and 2012 were sequenced in this study. Together with 5 isolates from 1997-2002 and 78 isolates collected from 2013-2017 sequenced previously [13], a total of 385 isolates were included in the analysis. Comparing to Tohama I, the number of SNPs identified per genome ranged from 191 to 308. A phylogenetic tree was constructed using the maximum likelihood method based on 1,501 SNPs (**Figure 1**). Among these 385 isolates, 351 were classified into our previously identified SNP Cluster I [7] which was characterised by the *ptxP3* allele and six cluster specific SNPs (**Supplementary Table 2 & 3**). The isolates carrying *ptxP3* allele were further divided into two sub-lineages demarcated by different *fim3* alleles, with 249 isolates carrying *fim3A* allele and 85 isolates carrying *fim3B* allele. Note that some studies used *fim3-1* and *fim3-2* notation for *fim3* alleles which is equivalent to *fim3A* and *fim3B* respectively.

Apart from the *ptxP3* lineage, 21 out of 34 isolates carried the *fim3A\** allele clustered into a defined *ptxP1-fim3A\** lineage. Twelve isolates which did not belong to any cluster were distributed between the *ptxP1-fim3A\** lineage and the *ptxP3* lineage. In addition, there was one *ptxP1* isolate which shared little homology with the major *ptxP1* clades and was located at the root of the tree with a long branch (**Figure 1**).

Our previous studies [19] identified five lineages which were named as epidemic lineages (ELs). The five ELs (EL1 to EL5) previously described [13] remained distinctive with expanded number of isolates, while a new EL (EL6) containing 21 isolates was identified as a sister group with EL1, EL3 and EL5 (**Figure 1**). All ELs were within the *ptxP3-fim3A* lineage, and the division of the ELs were supported by bootstrap values of >80% and specific SNPs (**Supplementary Table 3**). It should be noted that these lineages may have been circulating before the recent epidemics and thus the naming does not imply that they arose during the epidemic.

Our previous study showed that Prn-negative isolates emerged at the start of the first epidemic in 2008 before becoming predominant by the end of the epidemic in 2012. Prn-negative isolates continued to be predominant through the latest epidemic [3,13]. EL1 had the highest proportion of Prn-negative isolates which accounted for 96.7% (59/61), while EL5 and EL6 had the lowest proportion with 8.00% (2/25) and 9.52% (2/21), respectively. The proportion of Prn-negative isolates in EL2, EL3 and EL4 were 66.67% (12/18), 56.92 % (37/65) and 45% (18/40) respectively. The *ptxP3-fim3B* lineage also contained Prn-negative isolates with 27.05% (23/85) (**Figure 1**).

### Diversification of the epidemic lineages through the epidemics

Phylogenetic and BEAST analyses showed different ELs arose and diverged at different times. The root-to-tip phylogenetic distances of individual isolates showed a positive linear correlation with their collection dates (R^2^ = 0.1048, P = 6.481e-11) (**Supplementary Figure 1**), suggesting that *B. pertussis* evolved through successive SNP accumulation over time. The divergence time of *ptxP3* lineage was estimated to be around 1983 (95%CI, 1980 -1987).

To further understand how these lineages emerged, expanded or disappeared, as well as how the diversity of *B. pertussis* changed during the succession of lineages, lineage through time (LTT) analysis [32,36] was performed. As shown in **Figure 2A**, the diversifications [log (number of lineages)] of lineage *ptxP1-fim3A*, ptxP3-fim3B*, and *ptxP3-fim3A* started from around 1994, 1995, and 1997, respectively. Ten years after ACV replaced WCV in Australia, their diversifications peaked around 2007. In these three lineages, *ptxP1-fim3A\** failed to flourish during the 2008-2012 epidemic and declined rapidly, while diversification of both *ptxP3-fim3A* and *ptxP3-fim3B* accelerated during the epidemic. *B. pertussis* isolates carrying *ptxP3-fim3A* appeared later but also persisted longer. The diversification of the six ELs within the *ptxP3-fim3A* lineage were also assessed using LTT (**Figure 2B & C**). EL3, EL4 and EL5 started to diversify at similar time points. EL3 and EL4 persisted through the two epidemics while EL5 went extinct in the 2008-2012 epidemic. EL2 and EL6 started at similar times. EL2 declined gradually whereas EL6 disappeared abruptly. EL1 started the latest among the six ELs and declined gradually with the progression of the first epidemic. This genetic diversification analysis suggested that the pertussis epidemics from 2008 to 2017 consisted of overlapping expansion and decline of different lineages over time.

**Figure 2.**
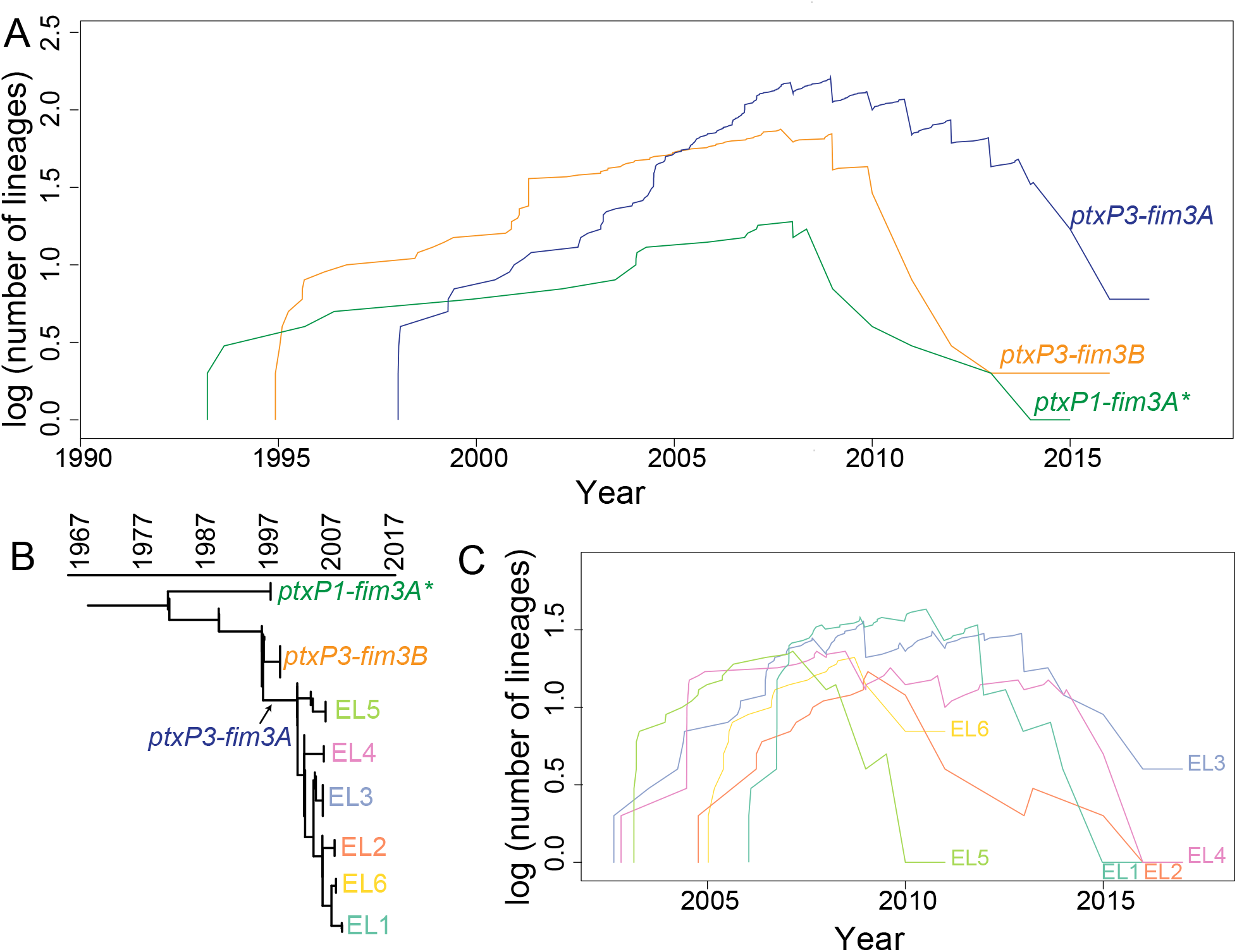
Lineage through time (LTT). LTT plot was based on the temporal distribution of nodes in the Australian *B. pertussis* phylogenetic tree. **A&C**: LTT for allelic types and the lineages identified from the phylogenetic tree. Y axis showed the log of the number of ancestral lineages; X axis showed the estimated emerging time of each lineage. Different lineages were coded by line colours. **B:** Schematic tree of Australian *B. pertussis* isolates showing phylogenetic relationships of allelic types and ELs.

### Patterns of intra-state and inter-state spread of *B. pertussis* in Australia

To examine how *B. pertussis* isolates spread across the country, we applied relative risk analysis [34] as a measure of transmission speed within defined intervals of divergence time and geographical distance using all 385 isolates based on the divergence date and tree in **Figure 1**. As shown in **Figure 3A**, two *B. pertussis* isolates collected from individuals residing within 1 km were five times more likely to share an MRCA of ≤ 1.5 years compared to two isolates from pairs which were > 260 km apart (the reference risk point). The probability of sharing an MRCA (<1.5 years) dropped to the reference point as the spatial distance increased between pairs and the value approached the reference risk (when distance > 20km). Furthermore, relative risks all approached the reference risk for the pair of isolates with a MRCA of ≥1.5 years (**Figure 3BCD**).

**Figure 3.**
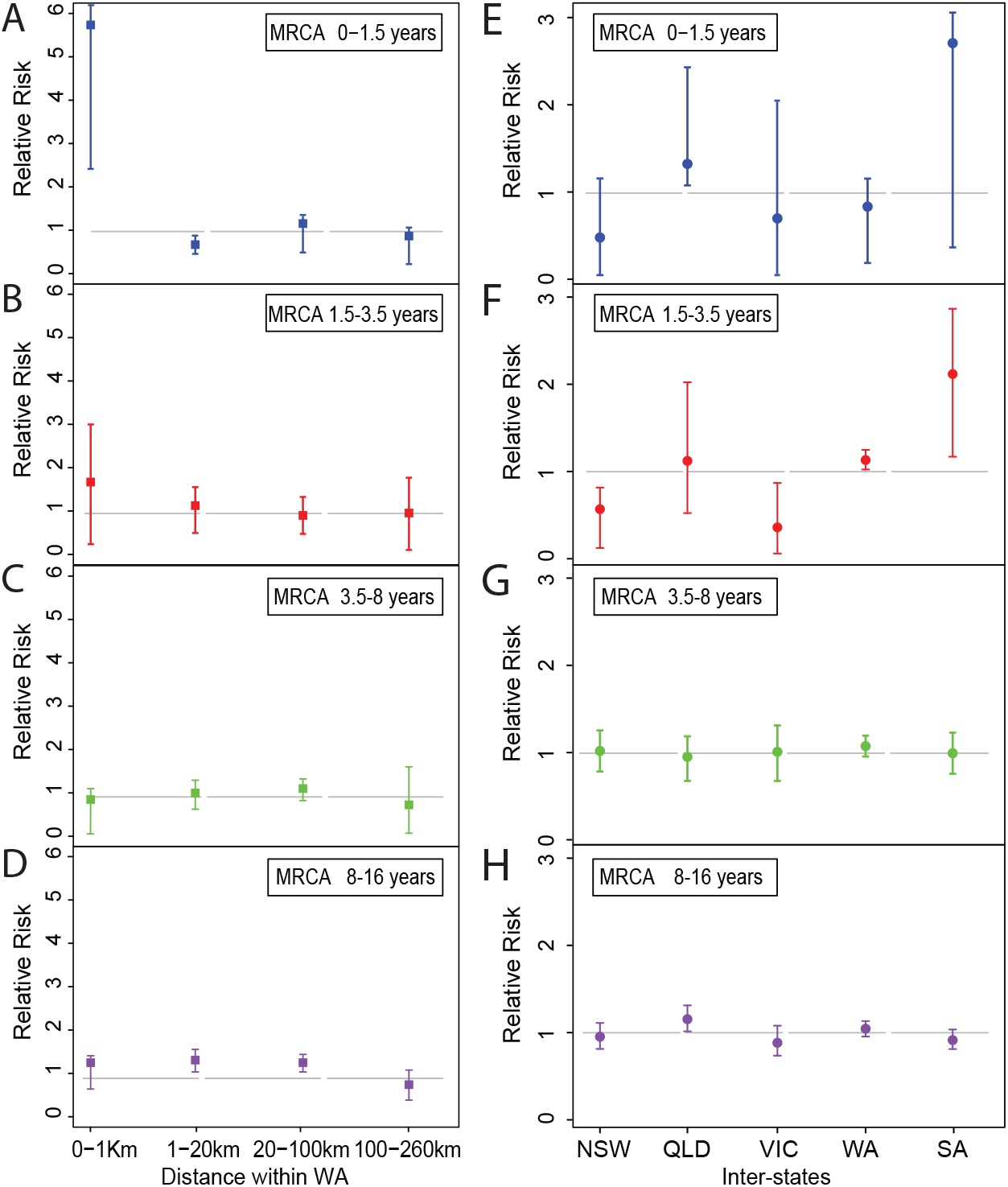
Relative risk analysis of Australian *B. pertussis* isolates within and between states. **A-D:** Intrastate WA (Western Australia) relative risk by MRCA within a defined period. Each point represents the risk that a pair of isolates collected from particular cases have an MRCA within a defined evolutionary timeframe relative to the risk that a pair of distal isolates (defined as two isolates from WA separated by > 260 km) have an MRCA in the same evolutionary timeframe. **E-H:** Interstate relative risk by MRCA within a defined period. Each point represents the risk that a pair of isolates collected from particular cases have an MRCA within a defined evolutionary timeframe relative to the risk that a pair of isolates from different state have a MRCA in the same evolutionary timeframe. One of the pair of isolates was selected from the specific state as shown in X-axis for each column and the other isolate was selected from any other 4 states. Each panel represents a different MRCA interval: A) MRCA < 1.5 years, B) MRCA 1.5 to 3.5 years, C) MRCA 3.5 to 8 years, D) MRCA 8 to 16 years. Grey broken lines represent the reference relative risks (reference ratio divided by itself, thus always equals one, see methods). Error bars represent the 95% CIs from 100 times bootstrap.

Relative risk was also used to assess the speed of *B. pertussis* isolates spreading across Australia. The analysis for relative risk of the <1.5 year pairs did not reach statistical significance at the 0.05 level possibly due to small number of isolates. When the MRCA period was >1.5 and < 3.5 years, it is clear that the relative risk of each state transmitting isolates to another state differed from the reference risk (**Figure 3E&F**). In contrast, the relative risk for each of the states where the isolates transmitted to another state approached the reference risk when the MRCA period was > 3.5 years (**Figure 3G&H**).

### Relationship of Australian epidemic lineages with global *B. pertussis*

A phylogenetic tree (**Figure 4**) of 1,452 *B. pertussis* genomes was constructed based on 5,777 SNPs in these genomes (See Methods), including 302 newly sequenced Australian *B. pertussis* isolates in this study and 1,150 publicly available genomes (**Supplementary Table 4**). Ten *ptxP* and *fim3* genotypes were identified with four major genotypes *ptxP1-fim3A/ptxP1-fim3A*, ptxP3-fim3A* and *ptxP3-fim3B* (**Figure 4**). These genotypes were phylogenetically clustered and the genotype divisions support the tree structure at large scale as expected, which is consistent with previous studies [4,37,38].

**Figure 4.**
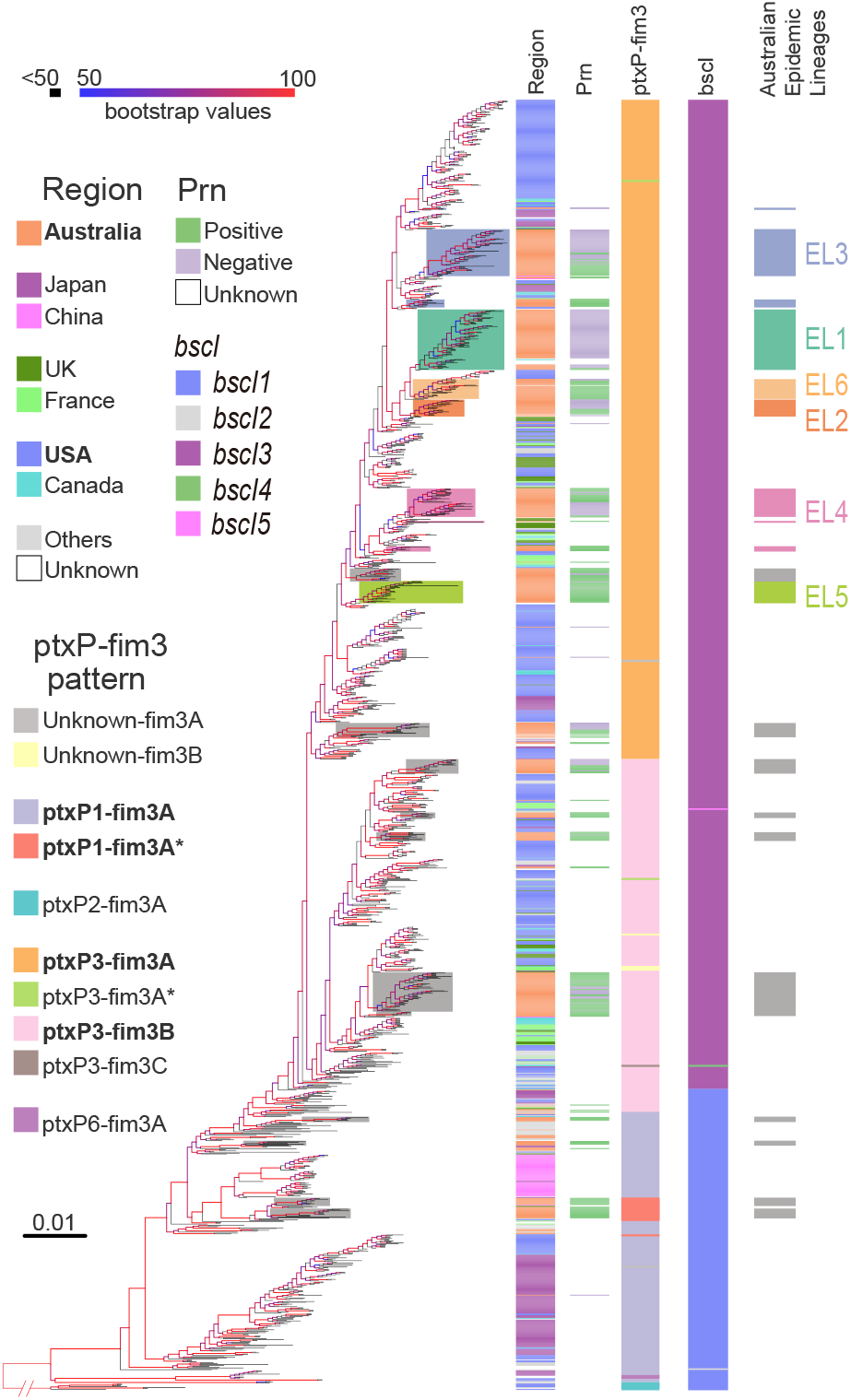
Phylogenetic tree of Australian and global *B. pertussis* isolates. Phylogenetic tree of 1,452 *B. pertussis* isolates based on genome-wide SNPs. The tree was rooted using a group of *ptxP2* isolates related to early Dutch strain B189 (isolated in 1991). The four columns to the right of the tree show the country of isolation, presence or absence of Prn, *ptxP-fim3* genotype and Australian lineages per colour legends. The background colours represent the ELs as indicated. For both tree branches and the 4^th^ Column, grey background colour of individual or groups of isolates marks Australian isolates not belonging to any ELs. Bootstrap values from 100 replicates are indicated by the colour of the branches as shown.

These isolates were collected from 1935 to 2017 across 20 countries in five continents, Asia, Africa, North and South America, Europe and Oceania (**Figure 4, Supplementary Table 4**). The largest number of isolates were from the US, Australia and Japan with 506, 404, and 180 strains respectively. Consistent with a previous study [37], the US isolates (annotated in blue in **Figure 4**) were distributed across the phylogenetic tree. Isolates from UK and France (annotated in dark/light green in **Figure 4**) were also generally distributed among *ptxP3-fim3A* and *ptxP3-fim3B* clades. In contrast, most (95%, 171/180) of the Japan *B. pertussis* isolates sampled between 1982-2013 (annotated in purple in **Figure 4**) were clustered in a single clade which was located near the root of the tree and were geographically restricted to Japan (**Figure 4**). Similarly, most (94%, 47/50) isolates from mainland China (annotated in pink in **Figure 4**) were grouped together in one clade.

The Australian isolates were spread across the tree with many instances of an Australian lineage consisting of one or more Australian isolates only sharing MCRA with isolates from another country (**Figure 4**), suggesting independent import into Australia. However, more than half of the isolates (230 of 404 Australian isolates grouped into the six ELs) were grouped together as distinctive lineages, in particular EL1, EL2, EL5 and EL6 (color-coded on branches in **Figure 4**). However, EL3 was split into 3 clusters with 1 large (52 isolates), 2 small clusters of 2 and 9 isolates, interspersed with isolates from other countries. Therefore, EL3 is not an exclusive Australian lineage and there were likely 3 independent origins of Australian isolates within EL3 with each cluster closest to different US isolates. Similarly, EL4 has also been split into 3 separate groups for Australian isolates with 1 large (32 isolates) and 1 small (6 isolates) clusters and 1 singleton. The large cluster was closest to UK isolates while the smaller cluster was closest to US isolates. Therefore, EL3 and EL4 are large global lineages with sub-lineages within the lineage being Australian only. By contrast, within EL1, EL2, EL5 and EL6, there were also isolates from other countries among the Australian isolates suggesting export of these lineages to other countries after expansion in Australia.

### Emergence and global expansion of *ptxP3/bscI3* genotype

Analysis of the phylogenetic tree (**Figure 4**) of 1,452 *B. pertussis* genomes revealed a major allelic replacement event in *bscI* (BP2249), which encodes the type III secretion system basal body protein BscI. From the 1,452 genomes, five *bscI* alleles (designated as *bscI1, bscI2, bscI3, bscI4* and *bscI5*) were observed. *bscI1* is the allele found in Tohama I. Relative to *bscI1, bscI2* has a non-synonymous SNP (G146A) which changes glycine to glutamic acid while *bscI3* has a non-synonymous SNP (A341G) which alters tyrosine to cysteine (**Figure 5A)**. *bscI4* and *bscI5* also contains the A341G mutation and an additional C184A and C131T mutation, respectively. C184A changes lysine to methionine in *bscI4* while C131T in *bscI5* changes alanine to valine (**Figure 5A**). A341G was predicted by PROVEAN to be a deleterious mutation while all other mutations were predicted to be neutral [20,39].

**Figure 5:**
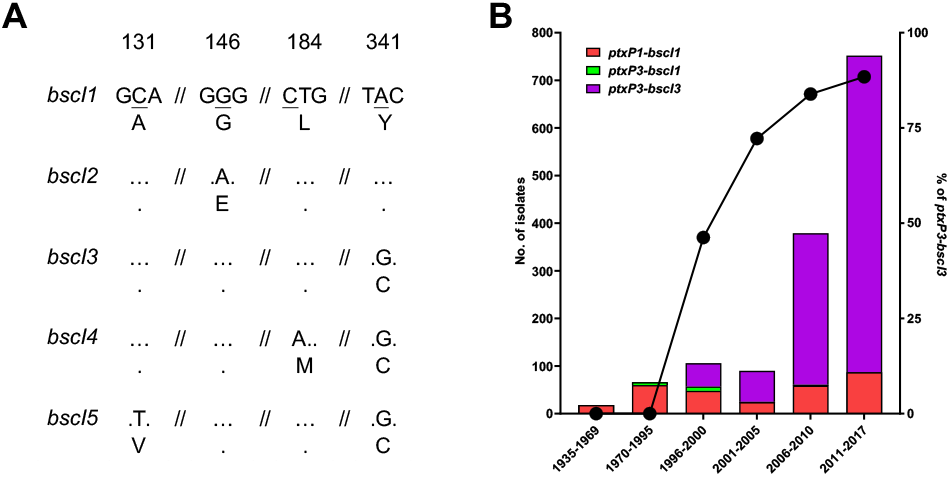
*bscI* alleles and their frequencies detected in 1,452 sequenced *B. pertussis* isolates from across the globe. (A) *bscI* alleles with nucleotide and amino acid changes. The numbers at the top refer to the position of the underlined SNP, relative to the start of the *bscI* gene. Dots represent identical base or amino acid. The nucleotide sequence is shown in codons with the corresponding amino acid (in single letter format) shown below. (B) Number and percentage of *ptxP3-bscI3* isolated since it was first detected in 1996. Red, green and purple bars depict the number of *ptxP1-bscI1, ptxP3-bscI1* and *ptxP3-bscI3* isolates. Twenty of the 1452 isolates were non-*ptxP1*/non-*ptxP3* or non-*bscI*1/non-*bscI3* and were not tallied in the graph. An additional 21 isolates had no collection date assigned and were also not tallied in the graph. The *ptxP1-bscI1, ptxP3-bscI1* and *ptxP3-bscI3* isolates were found across the globe. The line graph represents the rapid expansion of *ptxP3-bscI3* strains.

Of the 5 *bscI* alleles two, *bscI1* and *bscI3*, were predominant representing 23.3% (338/1452) and 76.5% (1111/1452) of the isolates, respectively. For *bscI2, bscI4* and *bscI5*, only 1 strain was identified for each. *bscI2* was detected in a *ptxP1-fim3A* strain while *bscI4* and *bscI5* were detected in a *ptxP3-fim3C* and *ptxP3-fim3B* strain, respectively. *bscI4* and *bscI5* appear to have evolved from a *bscI3* strain as both shared the *bscI3* mutation. The *bscI1*allele has been present since 1935 (the earliest strain in this study) and was mostly associated with *ptxP1* strains (**Figure 5B and Supplementary Table 5)**. As for *bscI3*, it appeared to have emerged as a single event (**Figure 4**) in a *ptxP3* strain with the earliest *bscI3* strain identified in 1996 from the Netherlands **(Supplementary Table 5)**. Between 1988-1995, all *ptxP3* strains sequenced had the *bscI1* allele, however after 1996, *ptxP3-bscI3* strains rapidly expanded and replaced the existing *ptxP3-bscI1* strains (**Figure 5B**). The last *ptxP3-bscI1* strain sequenced was isolated in 2009. From 2010 onwards, the *bscI1* allele has only been seen in strains carrying *ptxP1*, with many of the recent *ptxP1*-*bscI1* strains sequenced being isolated from China.

## Discussion

Australia experienced two major pertussis epidemics in the past two decades [13]. In a previous study by Safarchi *et al*. [19], a set of 27 isolates from the 2008-2012 epidemic was used to define five ELs, four of which expanded during the 2013-2017 epidemic [13]. This study expanded the previous studies with a larger dataset of 385 isolates, collected from 1997 to 2017, including isolates from the two recent epidemics (2008-2012 and 2013-2017) and compared with over 1000 global isolates. The Australian genomic data was also augmented by background epidemiological information, including isolation dates and locations, to allow comprehensive phylogeographic analyses to be performed and thus provided a more complete picture of *B. pertussis* population structure in Australia. This study has shown that some major lineages dominated during the two epidemics in Australia but there were also many independent lineages detected (**Figure 4**). These lineages had a shared MRCA with isolates from other countries and thus were likely to have been independently imported initially. Some lineages expanded while others died. Interestingly, the ELs were mostly imported as Prn-positive strains from other countries before expanding locally. These strains lost the Prn during the expansion in Australia, paralleling with the global reports of Prn-negative strains arising independently [11,40,41].

LTT analysis [32] depicted the development, disappearance and genetic diversification of different lineages across the two epidemics. The first epidemic (2008-2012) was associated with one *ptxP1* lineage and one *ptxP3* lineage. The outcompeting of *ptxP1* lineage by the *ptxP3* lineage started well before the epidemic and was consistent with the hypothesis of vaccine selection pressure [7,8]. The diversification of the *ptxP3* lineage began around 1995 to 1997 when ACV was first introduced in Australia [3], then peaked around 2008 at the start of the epidemic. During this period, two overlapping types of vaccine selective pressure co-existed: the waning but broader WCV-induced immunity and the newly introduced but narrower ACV immunity. In population genetics terms, above observations might be explained by hard and soft selective sweeps [42,43]. The emergence of the *ptxP3* lineage under vaccine selection led to hard selective sweeps across the population resulting in a population bottleneck, followed by rapid recovery of genetic diversity with multiple sub-lineages emerging within the *ptxP3* lineage. Further, the parallel emergence of Prn-negative strains from the selection pressure from the ACV can be viewed as soft selective sweeps followed by accelerated diversification of these sub-lineages. Therefore, hard and soft selective sweeps through vaccine selection pressures could have shaped the current *B. pertussis* population dynamics.

Over the course of the two epidemics, *ptxP3-fim3A* lineage gradually overtook *ptxP3-fim3B* lineage in Australia. Since Australia used mostly an ACV without Fim3 component, it is unlikely that ACV selection pressure played any role. There is a possibility that WCV played a role in the selection of strains in the early years of ACV era as vaccination or initial priming with WCV had a longer-term protection against *B. pertussis* infection than ACVs [44]. However, it is likely that, as the *B. pertussis* populations evolve, natural infection induced immunity may have affected the dynamics of spread of the two *ptxP3*-*fim3* sub-lineages. It is also possible that the two sub-lineages differed in frequency in their countries of origin, affecting their chances of being spread to other continents including Australia (the “founder effect”). In the US, *fim3B* was more prevalent in 2006-2009 while *fim3A* was more prevalent in the UK during 2001-2012 in the ACV era [45,46]. It is possible that strains with the *fim3B* allele were imported into Australia around 1995 (**Figure 4**) and expanded during the first epidemic. Another possible explanation could be the hitchhiking effect. When compared to *fim3B* isolates, a higher proportion of *fim3A* isolates have lost Prn which may have led to better survival of the *ptxP3-fim3A* lineage (**Figure 4**).

Further, LTT analysis of the six ELs within the *ptxP3-fim3A* lineage showed that the increase and decrease of the diversification of different ELs (EL1-6) occurred at different time points. The rise and fall of different ELs during the epidemic revealed that the pertussis epidemic was not due to a sole expansion of a single *ptxP3* lineage in Australia. Rather, in the epidemic period, the *B. pertussis* population was composed of different lineages with overlapping circulation of multiple lineages over time. Examination of the origins of the lineages in the global phylogenetic tree showed that four lineages, EL1, EL2, EL5 and EL6 were likely to have been imported into Australia once and then expanded as each of these lineages contained mostly Australian isolates. However, EL3 was now seen as consisting of one large (n=52) cluster and two smaller (n=2 and n=9) clusters of Australian only isolates in each case, suggesting that EL3 was a global lineage within which the clusters were actually imported into Australia and then underwent local expansion after importation (**Figure 4**). While EL4 has the same branching pattern. Within EL3, the large cluster expanded with emergence and expansion of a Prn-negative subcluster while the minor cluster contained no Prn-negative isolates (**Figure 4**). The data suggest that EL3 has adapted to circulate in Australia with the emergence of the Prn-negative subcluster. In EL4, Prn-positive isolates and Prn-negative isolates started to expand in parallel in two independent sub-lineages, but only Prn-negative sub-lineages survived after 2008. By contrast, EL6 which persisted during the first epidemic and then abruptly declined had only two Prn-negative isolates (9%). The above result suggests that the lineages with lower proportion of Prn-negative (e.g. EL6) isolates may not be fit enough to survive the competition for respiratory niches with other lineages with predominantly Prn-negative isolates. This can be interpreted as additional supporting evidence that the succession of the *B. pertussis* lineages was affected by the selection pressure from the vaccines.

Since isolates from this study had residential postcode data, the local spread of isolates was examined using relative risk analysis. Isolates that shared an MRCA within 1.5 years from present were more likely to be localised within 1 km (the approximate radius of an average suburb). In other words, once a new *B. pertussis* lineage emerged, it was more likely to spread locally within the first 1.5 years. However, any new lineage emerged after 1.5 years was likely to transmit to a wider region, reflecting rapid spread of pertussis strains within a state. Relative risk analysis was also used to examine the inter-state spread of *B. pertussis* in Australia. Within 3.5 years, newly arisen lineages were more likely to stay within a state. However, after 3.5 years, new lineages can spread between states with equal chances and these lineages were no longer highly spatially structured across the country.

Therefore, pertussis epidemics in Australia were characterised by the initial importation followed by expansions of different lineages over the epidemics. On the other hand, these lineages could also spread to other countries after a certain period of time, generating interweaving pertussis epidemics across the globe as seen with EL1 and EL6 within which isolates from other countries were found. Given that pertussis is a respiratory infection and is not confined by borders just like SARS-CoV-2 [47], epidemics in one country may rapidly affect other countries. However, the relative risk analysis suggests that strains were geographically restricted within and between states in the initial 1.5-3.5 years. Therefore, there was a likely lag of pertussis epidemics between countries from 1.5 to 3.5 years. These epidemic patterns are consistent with the time needed for a locally arisen lineage to spread.

Phylogenetic analysis of 1,452 Australian and international isolates revealed the emergence of *bscI3* in *ptxP3* strains. Although both *ptxP1*-*bscI1* and *ptxP3*-*bscI3* strains were circulating during the mid-1990s-2000s, the expansion of *ptxP3* strains in many ACV countries, such as Australia, US, UK, Canada and France, was only associated with the *bscI3* allele. This suggests that the *bscI3* allele may provide additional selective advantages to *ptxP3* strains. King *et al* [48] showed that *ptxP1* strains with a *ptxP3* genetic background had higher colonisation ability than wild type *ptxP1* strains, suggesting additional adaptation besides *ptxP3*. Our mouse studies found that *ptxP3* strains appeared to be fitter than *ptxP1* strains in both ACV immunised mice and naïve mice [23]. Further our previous proteomic studies suggests that the non-synonymous A341G mutation in *bscI3* may result in decreased secretion of T3SS effectors [20-22]. This may provide *ptxP3* strains with reduced immune recognition allowing it to evade the general host immune system and may partly explain the increased fitness of Cluster I strains regardless of immunisation status [23]. Since 2010, all *ptxP3* strains sequenced globally including *fim3A*/*fim3B* and Prn-positive/Prn-negative isolates were *ptxP3-bscI3* (with the exception of two *bscI4* and *bscI5* isolates which were derived from *bscI3*) suggesting a selective advantage of the A341G mutation. *ptxP3* strains were also found to predominant in Iran, a WCV country, which were all carrying *bscI3* allele [40]. There is a possibility that emergence of the *bscI3* allele was associated with WCV selection if BscI or the T3SS was part of the WCV antigens. Strains used to produce WCV most likely carried *bscI1* allele since all early circulation strains carrying *bscI1*. It is possible that *bscI3* was hitch-hiking on *ptxP3* but it remains to be seen whether the selective advantage provided by *bscI3* is dependent on the *ptxP3* allele. Most recent *ptxP1-bscI1* strains were currently circulating in China and its dominance may be associated with macrolide resistance [49].

## Conclusions

In this study, genome-scale phylogenetic analysis with spatial and temporal metadata revealed the important dynamics of importation and expansion of *B. pertussis* lineages in Australia. During the two epidemics in Australia, multiple *B. pertussis* ELs were initially imported, expanded and then declined at different time points. The changes were likely driven by multiple co-occurring forces including immune selection by vaccination and natural infection and transmission events. Our results also suggest that, once a new strain emerged, it was more likely to only spread within a short spatial distance in the first years after emergence. However, after 3.5 years of evolutionary time, the different lineages were no longer spatially structured across the country. This study showed that the expansion of *ptxP3* strains was also associated with the expansion of the *bscI3* allele. This allele may reduce secretion of T3SS effectors and allow *B. pertussis* to reduce immune recognition and thus increase overall fitness of *ptxP3* strains. These results suggest that the T3SS may be a good vaccine candidate. Together, these findings uncovered important characteristics of the epidemic dynamics and population structural changes of *B. pertussis* and improved our understanding of pertussis epidemics and re-emergence in highly vaccinated host populations.

## Acknowledgements

We thank Narelle Raven for technical assistance. This study was supported by a grant from the National Health and Medical Research Council of Australia (grant no.1146938). Z.X. was supported by a University of New South Wales scholarship.

## Data Availability

All Illumina sequencing reads analysed in this study were deposited in the NCBI database. Raw reads were deposited under the project accession number PRJNA562796. In addition, the other five Illumina sequencing reads collected from 1997-2002 were deposited under PRJNA280658 and 78 isolates from 2013-2017 Australian epidemic study were deposited under PRJNA432286. The global *B. pertussis* genomes were downloaded from Sequence Read Archive (SRA) in NCBI by searching “*Bordetella pertussis*” by the date of 6^th^ Nov 2018.

## Author Contributions

Ruiting Lan conceived the study, Zheng Xu, Dalong Hu and Laurence Luu performed the analysis. Sophie Octavia, Vitali Sintchenko, Frits R. Mooi and Laurence Don Wai Luu revised the manuscript. Mark M. Tanaka supported the calculation of the relative risk. Jenny Robson and Anthony D Keil collected *B. pertussis* isolates. Ruiting Lan provided critical analysis and discussions. All authors discussed the results and contributed to the revision of the final manuscript.

