## Supplementary_figure_1 for "Genomic dissection of the microevolution of Australian epidemic *Bordetella pertussis*"

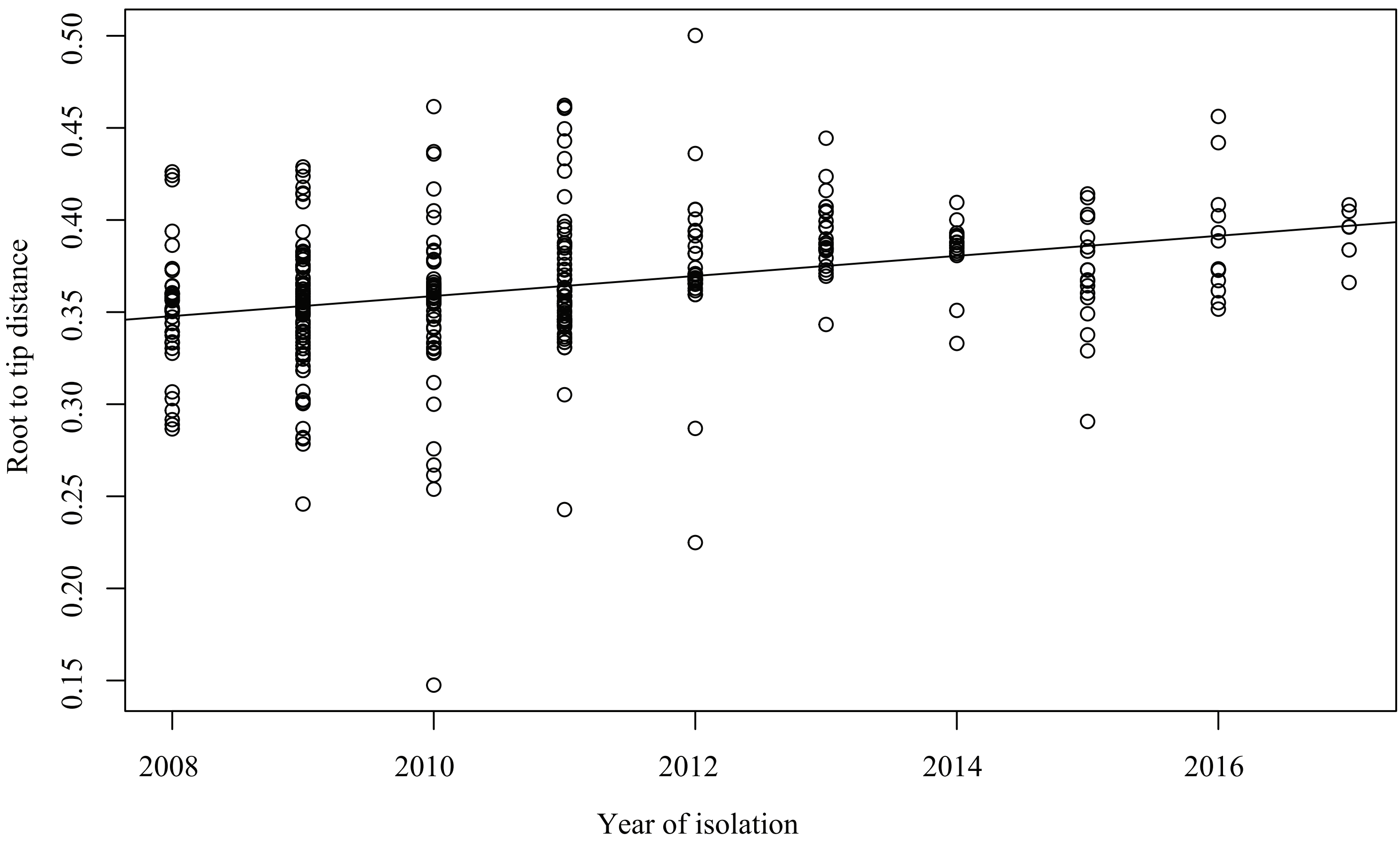


**Supplementary Figure 1.** Linear regression plot of *B. pertussis* displaying the correlation (R2) between the root-to-tips distance (y-axis) and the date of isolates (x-axis). The root-to-tip distances of individual isolates correlated with their date of isolation. (R2= 0.1048, P < 0.001)
